# Genetic perturbations of disease risk genes in mice capture transcriptomic signatures of late-onset Alzheimer’s disease

**DOI:** 10.1101/757161

**Authors:** Ravi S. Pandey, Leah Graham, Asli Uyar, Christoph Preuss, Gareth R. Howell, Gregory W. Carter

## Abstract

**Background:** New genetic and genomic resources have identified multiple genetic risk factors for late-onset Alzheimer’s disease (LOAD) and characterized this common dementia at the molecular level. Experimental studies in model organisms can validate these associations and elucidate the links between specific genetic factors and transcriptomic signatures. Animal models based on LOAD-associated genes can potentially connect common genetic variation with LOAD transcriptomes, thereby providing novel insights into basic biological mechanisms underlying the disease.

**Methods:** We performed RNA-Seq on whole brain samples from a panel of six-month-old female mice, each carrying one of the following mutations: homozygous deletions of *Apoe* and *Clu*; hemizygous deletions of *Bin1* and *Cd2ap*; and a transgenic *APOEε4*. Similar data from a transgenic *APP/PS1* model was included for comparison to early-onset variant effects. Weighted gene co-expression network analysis (WGCNA) was used to identify modules of correlated genes and each module was tested for differential expression by strain. We then compared mouse modules with human postmortem brain modules from the Accelerating Medicine’s Partnership for AD (AMP-AD) to determine the LOAD-related processes affected by each genetic risk factor.

**Results:** Mouse modules were significantly enriched in multiple AD-related processes, including immune response, inflammation, lipid processing, endocytosis, and synaptic cell function. WGCNA modules were significantly associated with *Apoe^−/−^, APOEε4, Clu^−/−^,* and *APP/PS1* mouse models. *Apoe^−/−^, GFAP-driven APOEε4,* and *APP/PS1* driven modules overlapped with AMP-AD inflammation and microglial modules; *Clu^−/−^* driven modules overlapped with synaptic modules; and *APP/PS1* modules separately overlapped with lipid-processing and metabolism modules.

**Conclusions:** This study of genetic mouse models provides a basis to dissect the role of AD risk genes in relevant AD pathologies. We determined that different genetic perturbations affect different molecular mechanisms comprising AD, and mapped specific effects to each risk gene. Our approach provides a platform for further exploration into the causes and progression of AD by assessing animal models at different ages and/or with different combinations of LOAD risk variants.

## BACKGROUND

Alzheimer’s disease (AD) is the most common adult neurodegenerative disorder and accounts for around 60-80% of all dementia cases [1]. Neuropathologically, Alzheimer’s disease is generally characterized by the presence of extracellular amyloid plaques composed of amyloid-β (Aβ) surrounded by dystrophic neurites, neurofibrillary tangles (NFTs), and neuronal loss [2, 3]. Clinically, AD is classified into two subtypes: early onset with Mendelian inheritance, and late onset (or sporadic) AD [1, 4]. Early-onset Alzheimer’s disease (EOAD) strikes prior to the age of 65 and accounts for approximately 5% of all AD cases, while the much more common late-onset Alzheimer’s disease (LOAD) is diagnosed at later life stages (> 65 years) [2, 5]. In comparison to rare casual variants in three genes: *amyloid precursor protein (APP), presenilin 1 (PSEN1),* and *presenilin 2 (PSEN2)* that contribute to EOAD [1, 6, 7], the genetics factors influencing LOAD are complex due to the interplay of genetic and environmental factors that influence disease onset, progression and severity [8, 9]. Before the era of large-scale genome wide association studies, the e4 allele of the *apolipoprotein E (APOE)* gene was the only well-established major risk factor for LOAD, accounting for about 30% of genetic variance [10, 11]. *APOEε4* was inferred to have moderate penetrance [11] with homozygous carriers having a roughly five-times-increased risk compared to those who inherit only one e4 allele of *APOE* [1, 12].

Identification of new AD-related genes is important for better understanding of the molecular mechanisms leading to neurodegeneration [7]. Genome-wide association studies (GWAS) have identified dozens of additional genetic risk loci for LOAD, with candidate genes including clusterin (*CLU*), bridging integrator 1 *(BIN1),* and CD2 associated protein *(CD2AP)* [1, 2, 7, 13]. These novel risk genes cluster in functional classes suggesting prominent roles in lipid processing, the immune system, and synaptic cell function such as endocytosis [1, 14]. Although these risk variants are often of small effect size, investigation of their functionality can reveal the biological basis of LOAD [1].

Despite recent advances in genetic and genomic resources to identify genetic risk factors, the disease mechanisms behind LOAD remain opaque. Most transgenic animal models are based on rare, early-onset AD genes which do not reflect the complete neuropathology or transcriptomic signatures of LOAD [15]. Although these transgenic mouse models were helpful to understand early molecular changes underlying Aβ and tau pathology, the corresponding genetic factors only account for a small fraction of AD. Thus, animal models based on LOAD-associated genes are necessary to connect common genetic variation with LOAD transcriptomes.

To better understand the molecular mechanism underlying LOAD, we performed transcriptome profiling and analyses from brain hemispheres of six month old female mice carrying mutations in LOAD-relevant genes *Apoe, Clu, Bin1,* and *Cd2ap.* Weighted gene co-expression network analysis identified several mouse modules significantly driven by *Apoe^−/−^* and *Clu^−/−^* mouse strains. Moreover, we have compared mouse modules with human postmortem brain modules from the Accelerating Medicine’s Partnership for AD (AMP-AD) to determine the AD relevance of risk genes. We observed enrichment of multiple AD-related pathways in these modules such as immune system, lipid metabolism, and neuronal system. This study of LOAD-relevant mice provides a basis to dissect the role of AD risk genes in AD pathologies.

## METHODS

### Mouse strains and data generation

All mouse strains were obtained from The Jackson Laboratory and maintained in 12/12-hour light/dark cycle (Table 1). All experiments were approved by the Animal Care and Use Committee at The Jackson Laboratory. RNA-Seq data were obtained from whole left hemisphere brain samples from a panel of six-month-old female mice carrying one of the following mutations in LOAD associated genes: homozygous deletion in *Apoe* and *Clu*; heterozygous deletion in *Cd2ap* and *Bin1;* and a transgenic *APOEε4* driven by a GFAP promoter on a *Apoe^−/−^* background (herein referred to as *Apoe^−/−^, Clu^−/−^, Cd2ap^+/−^, Bin1^+/−^* and *APOEε4*) (Table 1, [16–21]). There were six biological replicates for each late-onset model and control B6 mice. To minimize gene expression variation between mice, all mice in experimental cohorts were bred in the same mouse room and were aged together (to the extent possible). Cohorts were generated either by intercrossing heterozygous mice or in the case of *Bin1^+/−^* and *Cd2ap^+/−^* by crossing heterozygous mice to C57BL/6J (B6) mice, as homozygosity in these two genes is lethal. Data were also included from five whole left hemisphere brain samples from 6-month-old female mice from an early-onset AD model (APP/PS1, Table 1) [22] as well as seven additional B6 control replicates to account for batch effects.

**Table 1:**
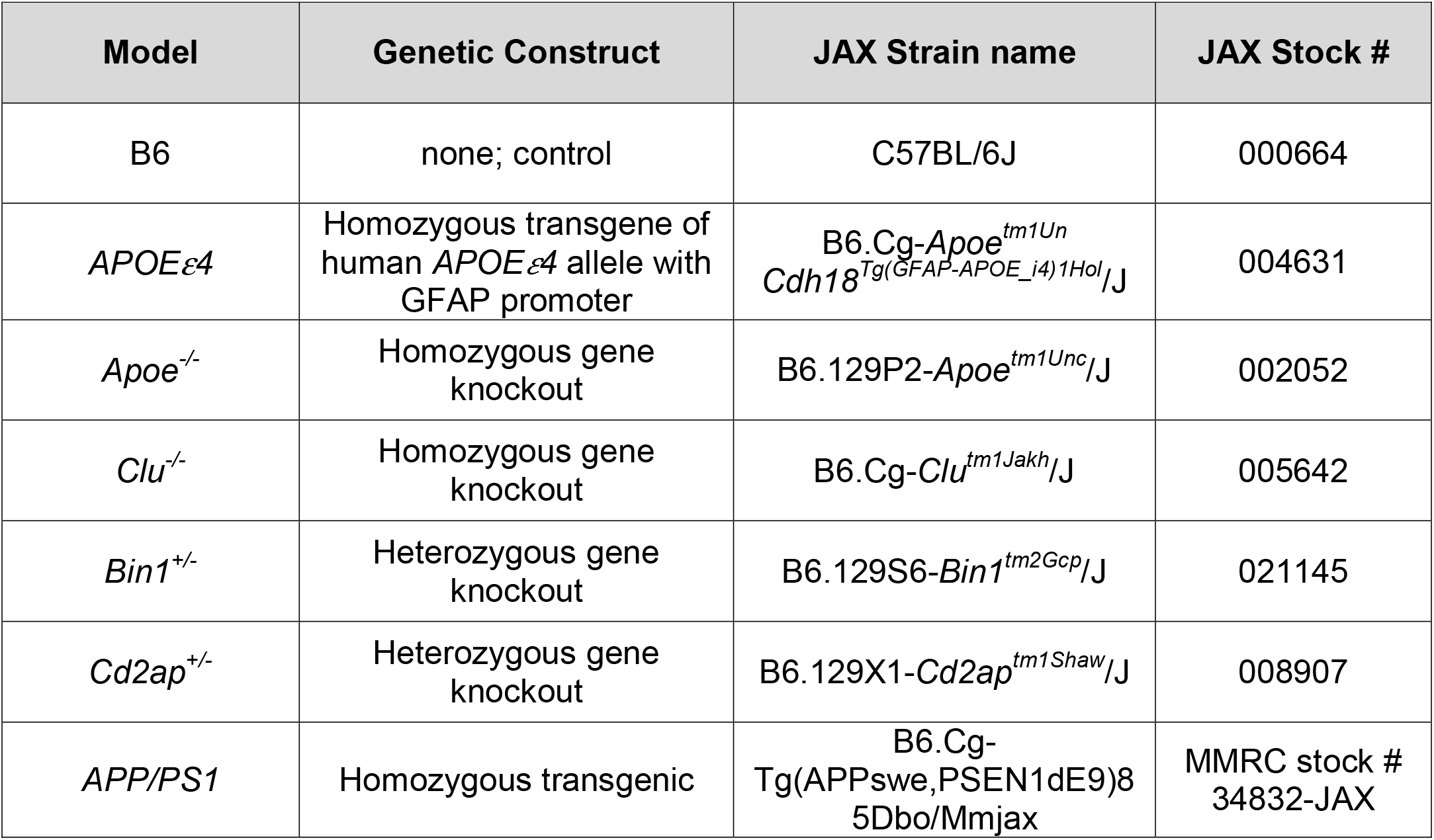
Study population. Whole-brain left hemispheres were collected at six months of age from female mice.

For sample collection, mice were anesthetized with a lethal dose of ketamine/xylazine, transcardially perfused with 1X phosphate buffered saline (PBS), brains carefully dissected and hemisected in the midsagittal plane. The left hemisphere was snap frozen. RNA extraction was performed using TRIzol (Invitrogen, cat #: 15596026) according to manufacturer’s instructions. Total RNA was purified from the aqueous layer using the QIAGEN miRNeasy mini extraction kit (QIAGEN) according to the manufacturer’s instructions. RNA quality was assessed with the Bioanalyzer 2100 (Agilent Technologies). Poly(A) selected RNA-Seq sequencing libraries were generated using the TruSeq RNA Sample preparation kit v2 (Illumina) and quantified using qPCR (Kapa Biosystems). Using Truseq V4 SBS chemistry, all libraries were processed for 125 base pair (bp) paired-end sequencing on the Illumina HiSeq 2000 platform according to the manufacturer’s instructions.

### Quality control of RNA-Seq data

Sequence quality of reads was assessed using FastQC (v0.11.3, Babraham). Low-quality bases were trimmed from sequencing reads using Trimmomatic (v0.33) [23]. After trimming, reads of length longer than 36 bases were retained. The average quality score was greater than 30 at each base position and sequencing depth were in range of 35 – 40 million reads.

### Read alignments and gene expression

All RNA-Seq samples were mapped to the mouse genome (assembly 38) using ultrafast RNA-Seq aligner STAR (v2.5.3) [24]. First, a STAR index was built from mm 10 reference sequence (Ensembl Genome Reference Consortium, build 38) for alignment, then STAR aligner output coordinate-sorted BAM files for each sample was mapped to mouse genome using this index. Gene expression was quantified in two ways, to enable multiple analytical methods: transcripts per million (TPM) using RSEM (v1.2.31) [25], and raw read counts using HTSeq-count (v0.8.0) [26].

### Differential expression analysis

Differential expression in mouse models was assessed using Bioconductor package DESeq2 (v1.16.1). [27]. DESeq2 take raw read counts obtained from HTSeq-count as input and has its own normalization approach. The significance of differential expression was determined by the Benjamini-Hochberg corrected p-values. The threshold for significance was set to corrected p=0.05. Differential expression in the *APP/PS1* models were obtained from a generalized linear model with the same significance threshold [22].

### Principal component analysis and batch correction

We analyzed 48 RNA-Seq samples originating from three experimental batches: 1) all late-onset genetic models (N=36); 2) one biological replicate of the *APP/PS1* strain with seven biological replicates of B6 control mice (N=8); and 3) four additional biological replicates of *APP/PS1* (N=4). First, we filtered out genes with TPM less than 10 for more than 90 percent of samples and then log-transformed to log2(TPM+1) for downstream analysis. We then used the plotPCA function of Bioconductor package EDASeq [28] to observe the differences in distribution of samples due to batch effects. Finally, we implemented COMBAT [29] on above RNA-Seq datasets to remove known batch effects.

### Network construction and mouse module detection

Modules (clusters) of correlated genes were identified using Weighted gene coexpression network analysis (WGCNA) implemented in R [30]. We used the step-by-step construction approach for network construction and module identification, which allows customization and alternate methods. The default unsigned network type was used, and a soft thresholding power of 8 was chosen to meet the scale-free topology criterion in the pickSoftThreshold function [31]. For module identification, WGCNA uses a topological overlap measure to compute network interconnectedness in conjunction with average linkage hierarchical clustering method. Modules correspond to branches of resulting clustering and are identified by cutting branches using dynamic tree cutting. To avoid small modules and ensure separation, we set the minimum module size to 30 genes and the minimum height for merging modules to 0.25. Each module is represented by the module eigengene (ME), defined as first principal component of the gene expression profiles of each module. Further, we have carried out one-way ANOVA (R function: aov) tests to determine differential expression between strains for each module eigengene. Modules with significant (p < 0.05) strain differences were analyzed for contributing strains using Tukey HSD (Tukey Honest Significant Differences, R function: TukeyHSD) for multiple pairwise-comparison between group means.

### Functional enrichment analysis

Functional annotations and enrichment analysis were performed using the R package clusterProfiler [32]. Gene Ontology terms and KEGG pathways enrichment analysis were performed using functions enrichGO and enrichKEGG, respectively, from the clusterProfiler package. The function compareCluster from this package was used to compare enriched functional categories of each gene module. The significance threshold for all enrichment analyses was set to 0.05 using Benjamini-Hochberg corrected p-values.

### Calculation and significance of Jaccard indices

Jaccard indices were computed to find overlap strengths between mouse modules and AMP-AD human modules. The Jaccard index is measure of similarity between sample sets and defined as ratio of size of the intersection to the size of the union of two sample sets. Further, to test the significance of the Jaccard index for each pair of mouse-human module overlap, we performed permutation analysis by random sampling the equivalent number of genes in each mouse module from the union of all genes in the mouse modules. This was performed 10,000 times to generate null distributions of Jaccard index values. Cumulative p-values were then calculated empirically.

### Transcription factor analyses

Transcription factors in mouse module were identified using iRegulon (v1.3) [33] in Cytoscape (v3.2.0) [34] and the Enrichr webtool that contains ENCODE and ChEA consensus transcription factor annotations from Chip-X library [35].

### Human post-mortem brain cohorts and co-expression module identification

Whole-transcriptome data for human post-mortem brain tissue was obtained from the Accelerating Medicines Partnership for Alzheimer Disease-(AMP-AD) consortium, which is a multi-cohort effort to harmonize genomics data from human LOAD patients. Harmonized co-expression modules from the AMP-AD data sets were obtained from Synapse (DOI: 10.7303/syn11932957.1). The human co-expression modules derive from three independent LOAD cohorts, including 700 samples from the ROS/MAP cohort, 300 samples from the Mount Sinai Brain bank and 270 samples from the Mayo cohort. A detailed description on post-mortem brain sample collection, tissue and RNA preparation, sequencing, and sample QC has been provided elsewhere [36–38]. As part of a transcriptome-wide meta-analysis to decipher the molecular architecture of LOAD, 30 co-expression modules from seven different brain regions across the three cohorts have been recently identified [39]. Briefly, Logsdon et al. identified 2,978 co-expression modules using multiple techniques across the different regions after adjusting for covariables and accounting for batch effects (10.7303/syn10309369.1). A total of 660 coexpression modules were selected based on a specific enrichment in LOAD cases when compared to controls (10.7303/syn11914606). Finally, multiple co-expression module algorithms were used to identify a set of 30 aggregate modules that were replicated by the independent methods [39].

### Correlation analysis

Standard gene set overlap tests are quick and easy, but do not account for direction of gene expression changes or coherence of changes across all genes in a module. To assess the directionality of genetic variants in model mice, we have computed the Pearson correlation across all genes in a given AMP-AD modules to determine human-mouse concordance.

To determine the effects of each genetic variant, we fit a multivariate regression model as:

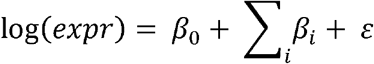

Where *i* denotes the genetic variants (*Apoe^−/−^, APOEε4, APP/PS1, Bin1^+/−^, Cd2ap^+/−^*, and *Clu^−/−^*), and *expr* represents gene expression measured by RNA-Seq transcripts per million (TPM).

Finally, we have computed the correlation (using cor.test function in R) between control versus AD log fold change (log2FC) in human derived modules and mouse gene expression effect (β) for the genes within an AMP-AD module. Log_2_FC values for human transcripts were obtained via the AMP-AD knowledge portal (https://www.synapse.org/#!Synapse:syn11180450).

## RESULTS

### Expression of target genes was modified by genetic perturbations

First, we have examined the relative expression (compared to control B6 mice) of LOAD associated genes to validate each strain. Expression of the mouse *Apoe* gene was downregulated in *Apoe^−/−^* mice (p < 1.00 × 10^−60^) as well as in transgenic *APOEε4* (p < 1.00 × 10^−258^) mice, which harbor human *APOE4* transcript driven by the GFAP promotor (Figure 1A). Expression of *Clu* gene was also downregulated (p < 1.00 × 10^−30^) in *Clu^−/−^* mice, while change in the expression of *Bin1* was significant but very small (log_2_FC = −0.3; p = 8.72 × 10^−12^) in *Bin1^+/−^* mice (Figure 1A). The change in expression of *Cd2ap* gene was not significant (log2FC= −0.07; p > 0.05) in *Cd2ap^+/−^* mice (Figure 1A). Overall, in each mouse strain, we observed significant downregulation in the expression of respective LOAD associated gene except in *Cd2ap^+/−^* models.

**Figure 1:**
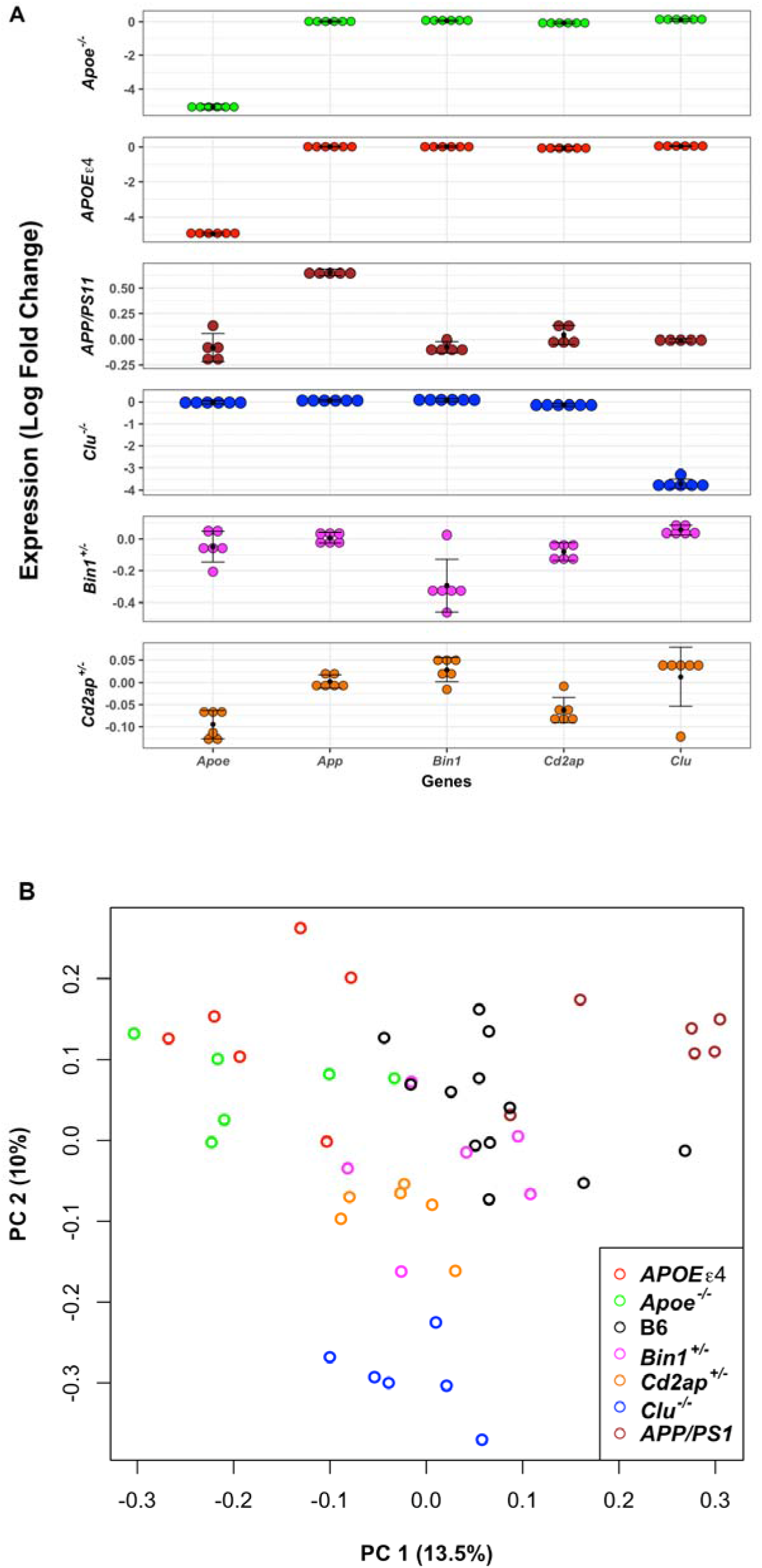
Expression of LOAD associated genes in mice. **(A)** Expression of AD associated risk genes in LOAD-relevant mice and the *APP/PS1* transgenic model compared to B6 (control) mice. X-axis shows AD-associated risk genes and Y-axis represents average log fold change expression of above genes in genetically perturbed mice compare to controls. **(B)** Principal component analysis of batch corrected RNA-seq data from mouse strains. The *APOEε4* (red circle) and *Apoe* KO (green circle) samples are most similar to each other. Samples from mice carrying only one copy of either *Bin1* (magenta circle) or *Cd2ap* (orange circle) occupy similar regions, which might be due to their related functions. *APP/PS1* samples (brown circle) were separated from mice with late-onset perturbations by the first PC.

### Transcriptional signatures from mice carrying different mutations in LOAD-relevant genes clustered into different groups by PCA

Principal component analysis (PCA) was performed on batch-corrected, log-transformed, and mean-centered TPM for 10,704 genes (Methods). The first principal component accounted for 13% of total variance and separated models of different types of AD: LOAD associated models and EOAD associated *APP/PS1* transgenic models cluster separately (Figure 1B), and thus might be affecting different AD-related processes. In other hand, within LOAD associated models, samples from the *Clu^−/−^* mice grouped together and separately from all other LOAD associated models in the second principal component (10% of variance) (Figure 1B). Across all strains, *APOEε4* transgenic and *Apoe^−/−^* mice were most similar to each other (Figure 1B). Hemizygous *Bin1^+/−^*, and *Cd2ap^+/−^* mice grouped closely to each another, suggesting functional similarity, and were the mutant strains in closest proximity to control (B6) mice (Figure 1B).

### Pathway analysis of differentially expressed genes identifies enrichment of different LOAD-related pathways in each mouse model

A total of 125 genes were significantly differentially expressed (p < 0.05) in *APOEε4* transgenic mice, out of which 55 genes were upregulated and 74 genes were downregulated (Table 2; Supplemental Table 1). We did not observe any pathway enrichment for differentially expressed genes in *APOEε4* transgenic mice. In *Apoe^−/−^* mice, 202 genes were identified significantly differentially expressed (p < 0.05), 137 genes were upregulated and 65 genes were downregulated (Table 2; Supplemental Table 1). Inflammation/immune response related pathways were enriched in the upregulated list of DE genes in *Apoe^−/−^* mice (Supplemental Table 2), as well as osteoclast differentiation that is related to *TREM2* and *TYROBP.* We did not observe any enrichment for downregulated genes in *Apoe^−/−^* mice. In *Clu^−/−^* mice, a total of 1750 genes were identified significantly differentially expressed (774 genes were upregulated and 976 genes were downregulated) (p < 0.05; Table2; Supplemental Table 1). Pathway analysis of DE genes identified endocytosis, RNA transport, and ubiquitin mediated proteolysis as enriched pathways in downregulated genes of *Clu^−/−^* mice, while ABC transporters, mTOR, and notch signaling as top enriched pathways in upregulated genes of *Clu^−/−^* mice (Supplemental Table 2). Only 20 and 36 genes were significantly differentially expressed (p < 0.05) in *Bin1^+/−^* and *Cd2ap^+/−^* mice, respectively (Table 2; Supplemental Table 1). Pathway analysis identified endocytosis, allograft rejection, autoimmune, type I diabetes as enriched pathways in downregulated genes of *Cd2ap^+/−^* mice (Supplemental Table 2), while there was no pathway enrichment in upregulated genes of *Cd2ap^+/−^* mice. Downregulated genes of *Bin1^+/−^* mice were enriched in signaling and FC gamma R-mediated phagocytosis pathways (Supplemental Table 2). In the *APP/PS1* transgenic mice, 624 genes were differentially expressed (303 and 321 genes were up and downregulated, respectively) (Table 2). Pathway analysis of these DE genes identified ribosome, oxidative phosphorylation, and Alzheimer’s disease as significantly enriched pathways (Supplemental Table 2).

**Table 2:**
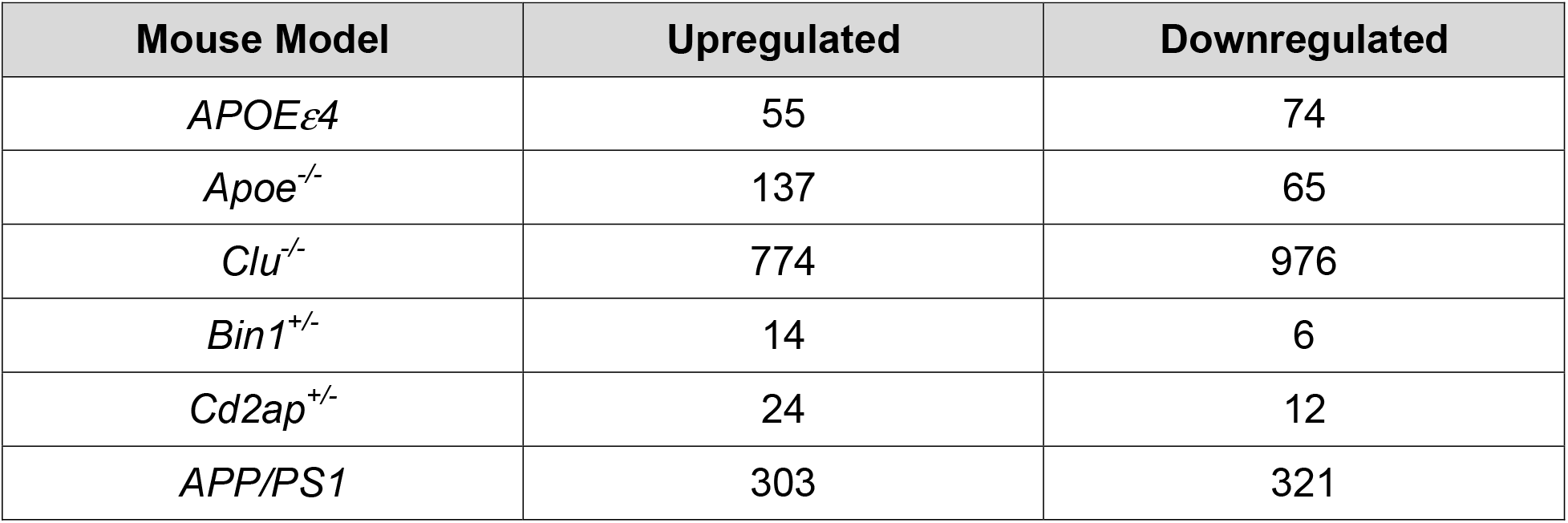
Differentially expressed genes by strain. Number of differentially expressed genes identified in each mouse strain compared to control mice (B6).

### Co-expression network analysis identified mouse modules enriched for multiple LOAD-related pathways driven by *APOE* and *CLU* strains

Weighted gene co-expression network analysis (WGCNA) [30] identified 26 distinct modules of co-expressed genes (Figure 2 A, Supplemental Table 3). Further, we have carried out one-way ANOVA test followed by Tukey-HSD (see methods) to determine if there was differential expression between strains for each module eigengene. We identified that 13 out of 26 modules were significantly driven by one or more of *Apoe^−/−^, APOEε4, Clu^−/−^*, and *APP/PS1* models (Supplemental Table 3). Pathway enrichment analysis identified that multiple AD-related pathways were significantly enriched in these mouse modules. *Apoe^−/−^* mice were significantly associated with ivory module (N = 64, p = 9.7 × 10^−6^), while the skyblue3 (N = 80, p = 4.6 × 10^−13^) (Figure 3; Figure 4; Supplemental Table 3) module were significantly associated with both *Apoe^−/−^* and *APOEε4* strains. Pathway analysis identified that the ivory mouse module was enriched in inflammation and microglia related pathways such as osteoclast differentiation, staphylococcus aures infection, phagosome, and endocytosis (Figure 2B), implicating an important role of *Apoe* in inflammatory and microglia related functions [40–42]. Brown (N = 1778, p = 3.1 × 10^−7^), lightcyan1 (N = 1206, p = 1.9 × 10^−5^), black (N = 685, p = 2.0 × 10^−2^), plum1 (N = 80, p = 1.0 × 10^−2^), and brown4 (N = 55, p = 0.04) modules were significantly associated with *Clu^−/−^* (Figure 3; Figure 4; Supplemental Table 3). The steelblue module was driven by both *Clu^−/−^* (p = 5.02 × 10^−13^) and *Cd2ap^+/−^* models (p = 9.5 × 10^−13^) (Figure 3; Figure 4; Supplemental Table 3). These mouse modules were enriched in many different pathways particularly related to synaptic cell function, endocytosis, and RNA transport (Figure 2B). This suggest the role of *Clu* gene in synaptic/neuronal related functions, which is in consistent with findings that reduced expression of *Clu* may results to aberrant synaptic development and neurodegeneration [43]. The darkorange2 (N = 61, p = 1.0 × 10^−6^), darkorange (N = 312, p = 0.03), orange (N = 142, p = 4.64 × 10^−13^), and lightgreen (N = 1456, p = 1.0 × 10^−12^) modules were found to be driven by *APP/PS1* (Figure 3; Figure 4; Supplemental Table 3). The lightyellow module (N = 163) was observed to be associated with both *APP/PS1* (p = 8.7 × 10^−5^) and *Clu^−/−^* mice (p = 1.4 × 10^−2^), but more significantly with *APP/PS1* (Figure 3; Figure 4; Supplemental Table 3). *APP/PS1*-driven modules (lightyellow, lightgreen, darkorange2) were enriched in lipid-processing and metabolism related pathways (Figure 2B). None of the modules were observed to be associated with *Bin1^+/−^* and *Cd2ap^+/−^* mice alone.

**Figure 2:**
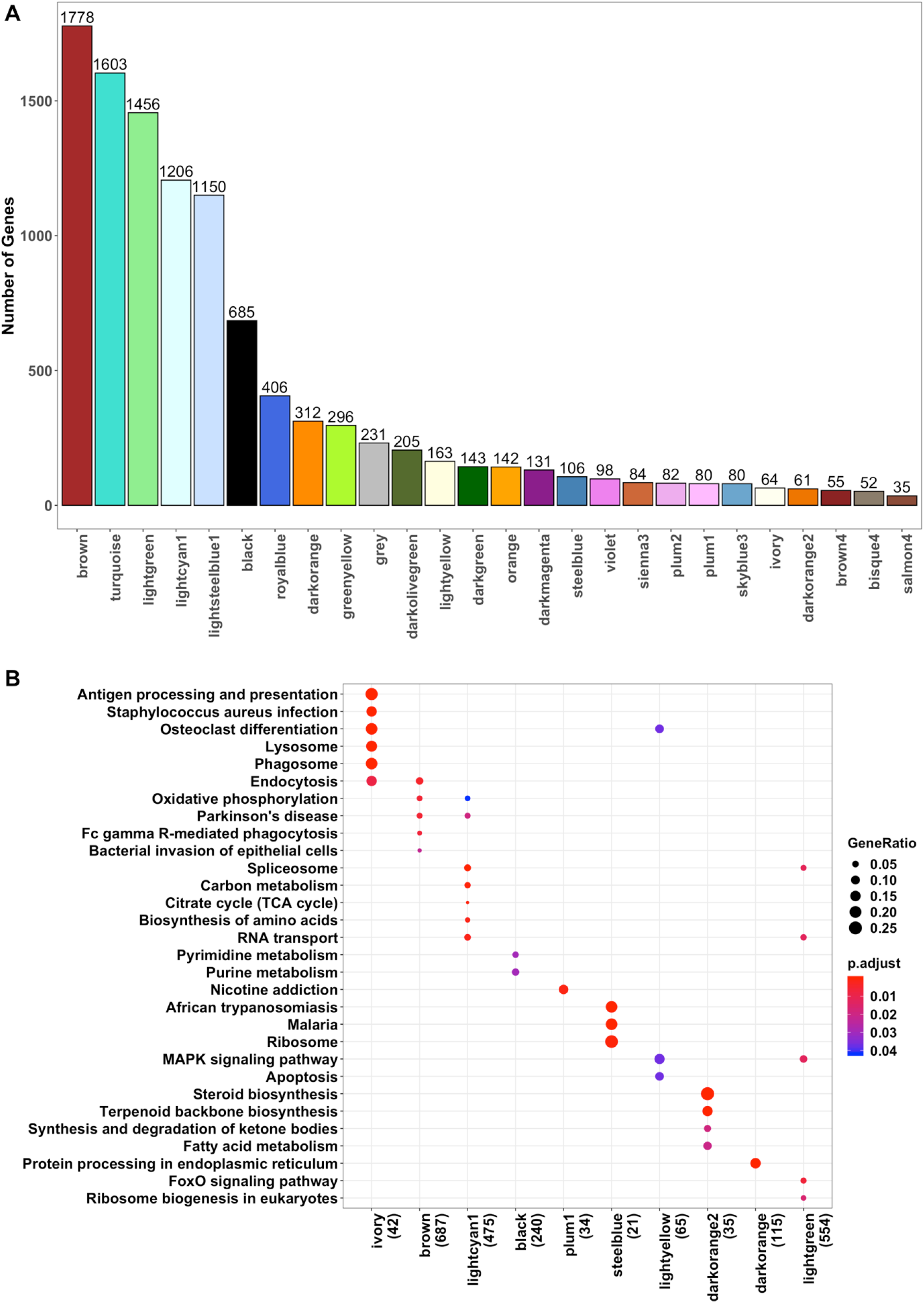
Mouse Modules Identified through WGCNA. **(A)** Twenty-six distinct mouse modules were identified from 10,704 mouse genes using WGCNA. Mouse modules of various sizes represented by different color names. **(B)** KEGG Pathway enrichment analysis (p < 0.05) in mice using enrichKEGG function build under clusterprofiler R package.

**Figure 3:**
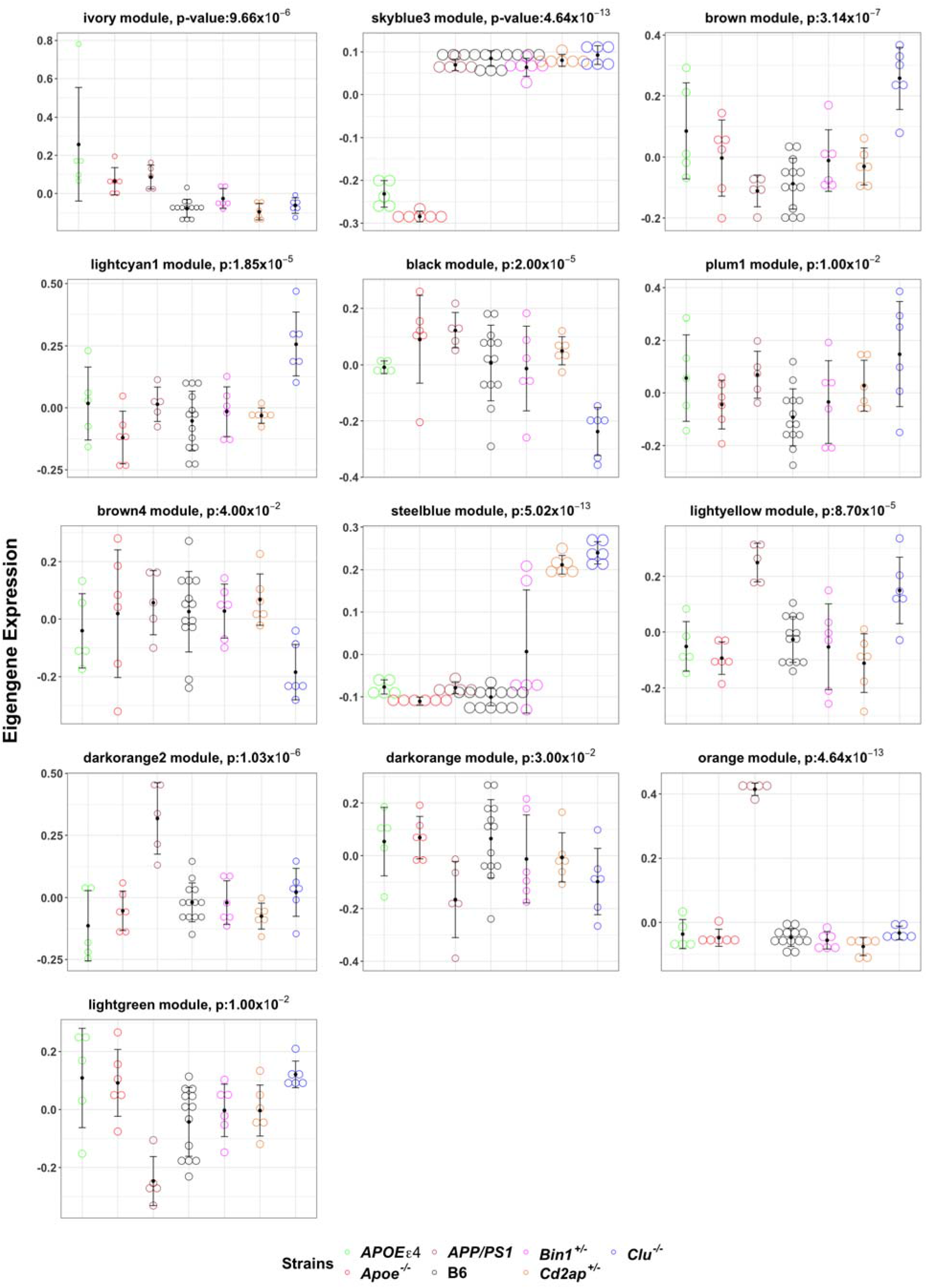
Mouse Modules Significantly driven by specific mouse strains. Expression of module eigengenes in mouse modules significantly driven by *Apoe^−/−^, APOEε4, Clu^−/−^* and *APP/PS1* mice (arbitrary units).

**Figure 4:**
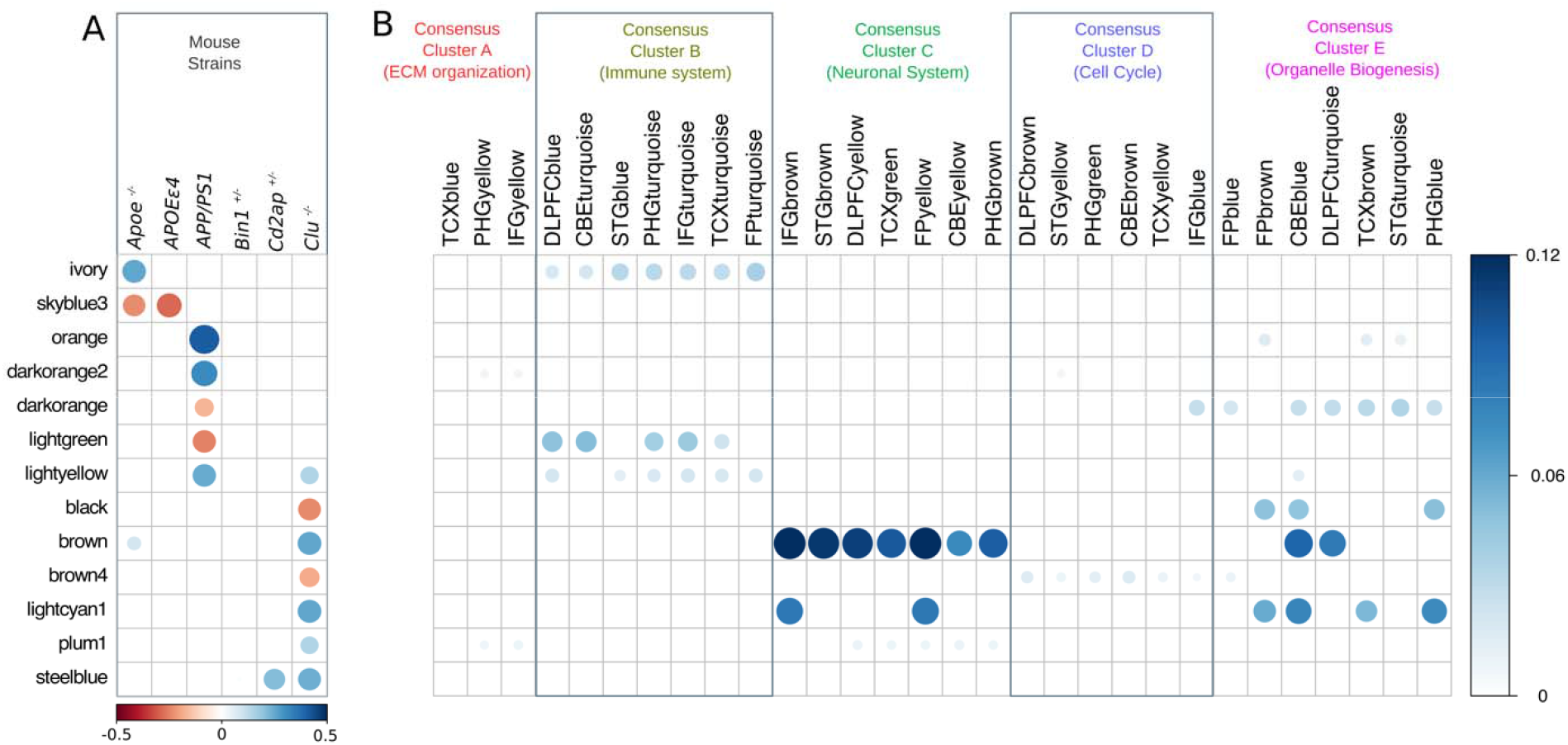
Overlaps between strain-associated mouse modules and human AMP-AD modules. **(A)** Mouse modules significantly driven by one or more of *Apoe^−/−^*, APOEε4, APP/PS1, Cd2ap^+/−^ and Clu^−/−^ mouse strains. The horizontal scale bar represents the average eigengene expression of mouse strains in mouse modules. **(B)** Overlaps between mouse modules and 30 human AMP-AD modules. The vertical scale bar represents Jaccard indices between mouse modules and AMP-AD modules. Jaccard indices were computed between each mouse and AMP-AD human modules.

### Comparison of mouse and AMP-AD modules

Finally, we compared mouse modules with the 30 human postmortem brain modules from the Accelerating Medicine’s Partnership for AD (AMP-AD). We computed Jaccard indices and its significance for each mouse – human module pair to identify which mouse module significantly overlap with human modules in order to identify AD-relevance of risk genes (Supplemental Table 5). Since each human module was derived from a specific brain region and study cohort, there are significant similarity between AMP-AD modules. Overlapping modules were therefore grouped into Consensus Clusters [39].

### *Apoe-*driven mouse module overlapped with AMP-AD inflammation and microglial consensus cluster

The ivory mouse module driven by *Apoe^−/−^* significantly overlapped with AMP-AD inflammation and microglia modules in Consensus Cluster B [39] (Figure 4; p < 0.05) and ranked among top ten mouse-human modules overlap (based on Jaccard indices) (Supplemental Table 4). These findings imply the significant role of *Apoe* in inflammation and microglia-related pathways. Furthermore, we identified that 22 genes were present in all AMP-AD microglial modules in Consensus Cluster B as well as in the Apoe^−/−^-driven ivory module (Figure 5), as these genes were expressed from all human brain regions and therefore might be playing the important role in inflammation and microglia associated pathways. In order to identify transcriptional changes in these genes due to any AD-relevance genetic alteration, we assessed differential expression of these 22 genes in each mouse model (Supplemental Table 1). Nine out of these 22 genes *(TREM2, CSF1R, C1QA, C1QB, C1QC, PTGS1, AIF1, LAPTM5* and *LY86)* were significantly upregulated (p < 0.05) in *Apoe^−/−^* mice and one gene *(TYROBP)* was significantly downregulated (p < 0.05) in *Clu^−/−^* mice. Some of these genes *(TREM2, TYROBP, C1QA,* and *CSF1R)* have been associated with AD and reported to be potential drug targets (https://agora.ampadportal.org/). We did not find a significant overlap between the skyblue3 mouse module and any AMP-AD module.

**Figure 5:**
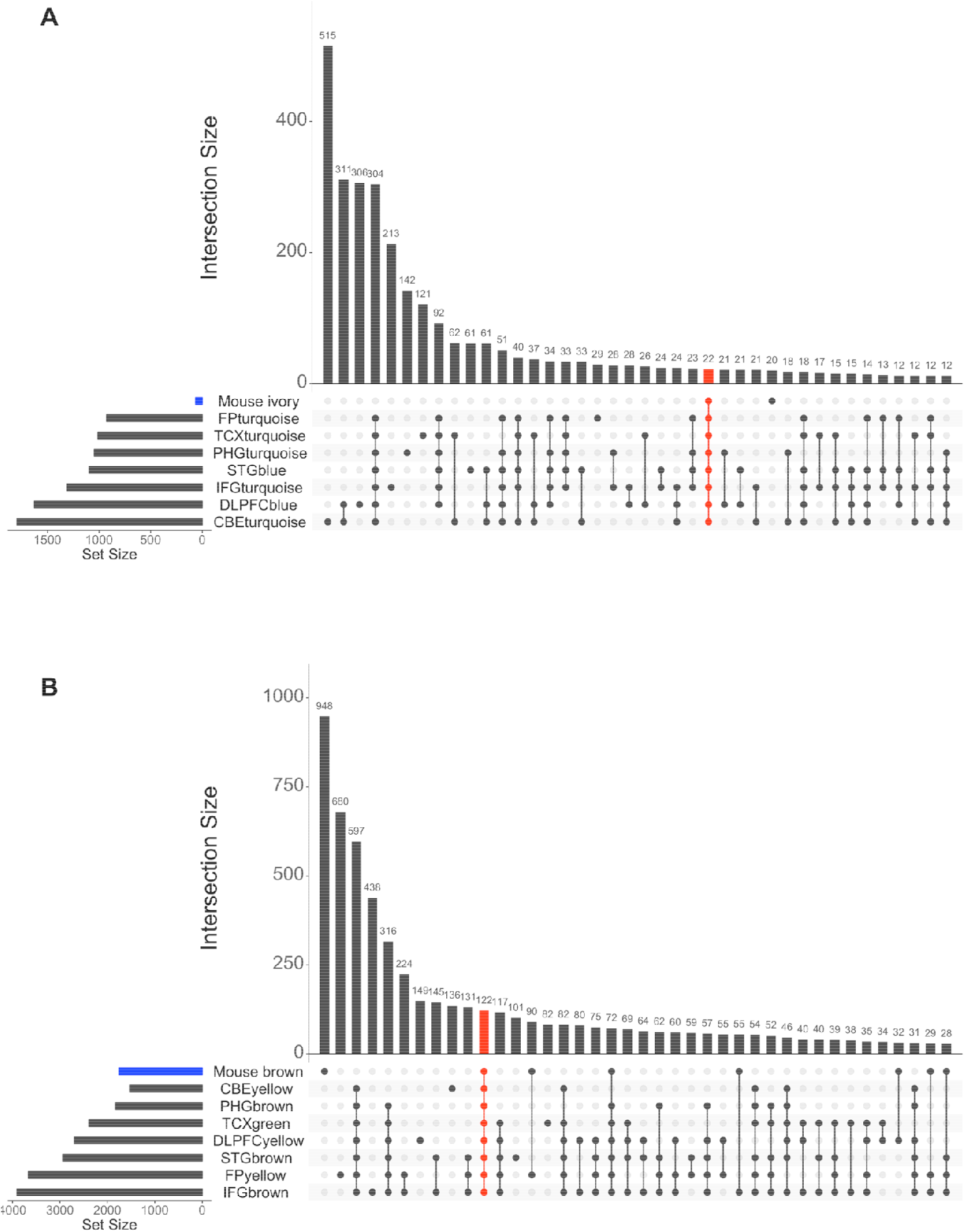
Overlaps between AMP-AD and key mouse modules: **(A)** Overlap between AMP-AD microglia modules in Consensus Cluster B and *Apoe^−/−^*-driven ivory module (shown in blue). We identified 22 genes which were present in all AMP-AD microglia modules in Consensus Cluster B and the mouse ivory module (red vertical bar). **(B)** Overlap between AMP-AD neuronal modules in Consensus Cluster C and *Clu^−/−^* driven brown module (shown in blue). We identified 122 genes which were present in all AMP-AD neuronal modules in Consensus Cluster C and mouse brown module (red vertical bar).

***Clu-driven* modules overlapped with AMP-AD neuronal system consensus cluster** *Clu^−/−^*-driven mouse modules (brown, lightcyan1, and plum1) prominently overlapped with AMP-AD neuronal system modules in Consensus Cluster C [39], while black, lightcyan1, and brown modules overlapped with organelle biogenesis associated AMP-AD modules in Consensus Cluster E (Figure 4; p < 0.05). The *Clu^−/−^*-driven brown4 module showed association with cell cycle associated AMP-AD modules in Consensus Cluster D (Figure 4; p < 0.05). Also, we have observed that the top five mouse-human module overlaps (based on Jaccard indices) were between the brown module and AMP-AD neuronal system modules in Consensus Cluster C (Supplemental Table 4). Further, we also identified that 122 genes were common between the *Clu^−/−^*-driven brown mouse module and all AMP-AD neuronal system modules in Consensus Cluster C (Figure 5B). We assessed these 122 genes for differential expression in each mouse strain (Supplemental Table 1) and found that 35 out of these 122 genes were differentially expressed (30 genes were upregulated and 5 genes were downregulated) only in *Clu^−/−^* mice, while three out of these 122 genes were differentially expressed only in *APP/PS1* transgenic mice (one gene was upregulated and two were downregulated). One of these 122 genes *(Syt7)* was upregulated in both *Clu^−/−^* mice and the *APP/PS1* transgenic mice. These finding suppot the likely role of *CLU* in neuronal function.

### *APP/PS1*-driven modules overlapped with inflammation, lipid-processing, and metabolism AMP-AD modules

The *APP/PS1*-driven orange and darkorange modules overlapped with lipid processing and metabolism associated AMP-AD modules in Consensus Cluster E, the lightgreen module overlapped with immune system modules Consensus Cluster B, and the lightyellow module overlapped with both microglia and organelle biogenesis related AMP-AD modules in Consensus Clusters B and E, respectively (Figure 4; p < 0.05). We found significant overlap for the darkorange2 mouse module with AMP-AD modules in Consensus Cluster E, which are in turn enriched in organelle biogenesis related pathways (Figure 4; p < 0.05).

### Correlation analysis provides directional coherence between mouse models and AMP-AD Consensus Clusters

The gene set overlap analysis identified mouse modules that are significantly overlapped with AMP-AD modules, but it does not assess directional coherence between AMP-AD modules and the effects of genetic perturbations in mice. To address this issue, we computed the Pearson correlation between log fold change gene expression in human AD cases versus controls (Log_2_FC) and the effect of each mouse perturbation on mouse orthologs as determined by the linear model (β) for the genes within an AMP-AD module. *Apoe^−/−^* and *APOEε4* mice showed significant positive correlation (r = 0.1 – 0.3, p < 0.05) with immune associated AMP-AD modules in Consensus Cluster B and significant negative correlation (r = −0.05, p < 0.05) with AMP-AD neuronal modules in Consensus Cluster C (Figure 6). Furthermore, *Clu^−/−^ and Cd2ap^+/−^* mice showed significantly positive association (r = 0.1, p < 0.05) with AMP-AD neuronal modules in Consensus Cluster C and negative correlation (r = −0.15, p < 0.05) with AMP-AD immune related modules in Consensus Cluster B (Figure 6). *Bin1^−/−^* and *APP/PS1* mice showed significant positive correlation (r = 0.1 – 0.2, p < 0.05) with AMP-AD immune response associated modules in Consensus Cluster B as well as AMP-AD neuronal modules in Consensus Cluster C. The cell cycle and RNA non-mediated decay pathways enriched AMP-AD modules in Consensus Cluster D were significantly negatively correlated (r = −0.2, p < 0.05) with *Apoe^−/−^, APOEε4, Clu^−/−^, Cd2ap^+/−^*, and *APP/PS1* mice, but *Bin1^+/−^* mice showed significant positive correlation (r = 0.11, p > 0.05) with AMP-AD cell cycle module in the cerebellum (Figure 6). Most of the AMP-AD modules in Consensus Cluster E that is enriched for organelle biogenesis associated pathways showed significant negative correlation (r = −0.1, p < 0.05) with all strains except the *Apoe^−/−^* models (r = 0.12, p < 0.05), while the AMP-AD modules of Consensus Cluster E in the frontal pole (FPbrown) and parahippocampal gyrus (PHGblue) showed significant positive association (r = 0.05 – 0.2, p < 0.05) with all strains (Figure 6).

**Figure 6:**
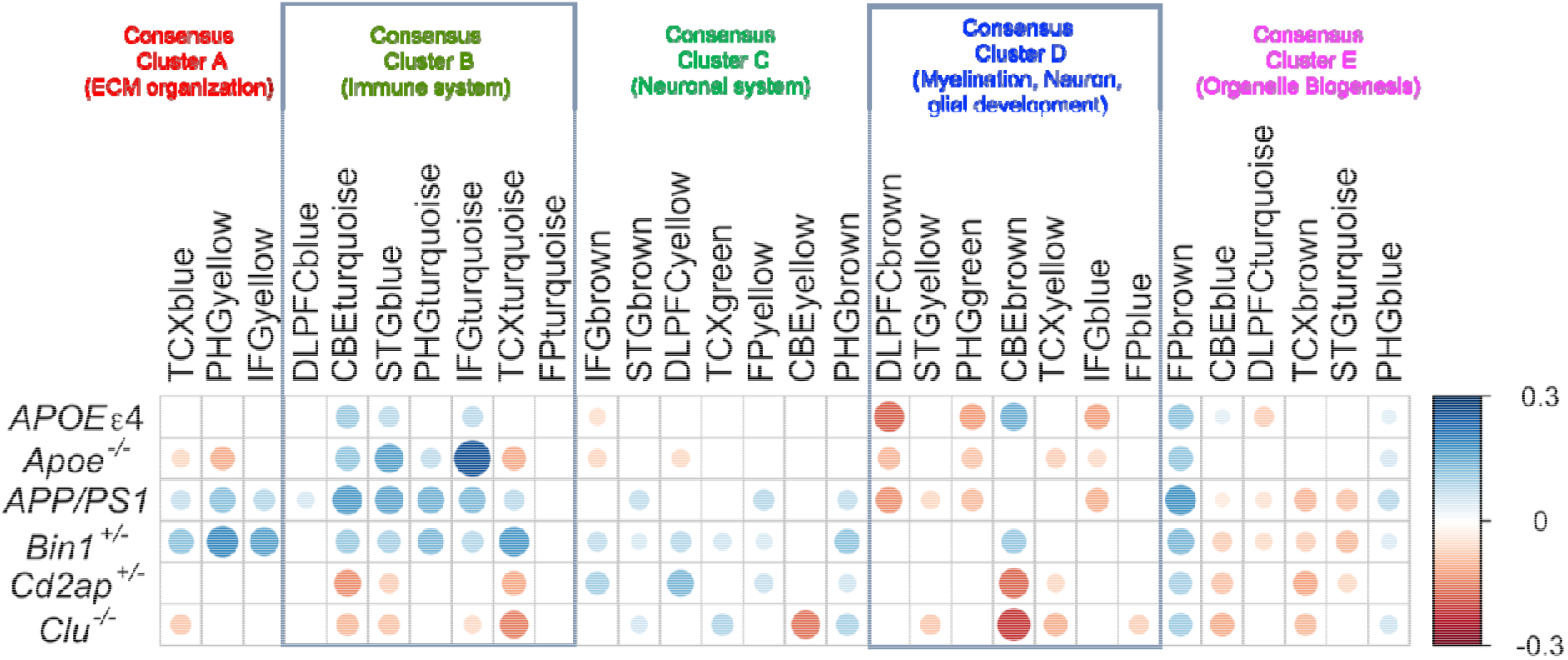
Correlation between mouse strains and 30 AMP-AD modules. Pearson correlation coefficients between 30 human AMP-AD modules and mouse strains. AMP-AD modules are grouped into five previously-identified consensus clusters describing the major functional groups of AD-related alterations. The vertical axis represents AMP-AD modules and the horizontal axis represents mouse strains. Positive correlations are shown in blue and negative correlations in red color. Color intensity and size of the circles are proportional to the correlation coefficient. Correlations with p-value > 0.05 are considered insignificant and are leaved blank.

### *Apoe-*associated modules are enriched in SPI1 regulatory targets

Transcription regulation play an important role in the initiation and progression of AD [44]. Our results provide evidence of the AD relevance of risk genes, but it is also important to identify the regulatory elements and transcriptional factors that regulate the expression of these genes for molecular dissection of disease etiology [45, 46]. Recent study have shown that *APOEε4* genotype suppress transcription of autophagy mRNA’s by competing with transcription factor EB for binding to coordinated lysosomal expression and regulation(CLEAR) DNA motifs [47]. TFs were identified for each module with high normalized enrichment scores (NES ≥ 4) from iRegulon (Methods), which correspond to an estimated false discovery rate of less than 0.01 [33] (Supplemental Table 5). The SPI1 transcription factor was enriched for regulatory targets in the *Apoe^−/−^* driven ivory and skyblue3 modules (Table S6). It has been previously reported that SPI1 responds to inflammatory signals and regulates genes that can contribute to neurodegeneration in AD [48]. We also observed that transcription factors from ELF, ETS, TCF, PEA3, GABP, and ERF sub-family of the E26 transformation-specific (ETS) family were enriched in the *Clu^−/−^*-driven modules (Supplemental Table 5). ETS-domain proteins play a role in the regulation of neuronal functions [49]. ETS family members ELK1 and ETS1 have been reported to expressed in neuronal cells and activate transcription of early onset AD candidate gene *PSEN1* [44, 50]. Understanding the role of these and other transcription factors in regulating AD associated genes can provide a molecular basis for potential therapeutic development.

## CONCLUSIONS

In this study, we have performed transcriptomic analysis of mouse strains carrying different mutations in genes linked to AD by GWAS to better understand the genetics and basic biological mechanisms underlying LOAD. We have also performed a comprehensive comparison at the transcriptomic level between mouse strains and human postmortem brain data from LOAD patients. This study of LOAD-relevant mouse models provides a basis to dissect the role of AD risk genes in relevant AD pathologies.

We determined that different genetic perturbations affect different molecular mechanisms underlying AD, and mapped specific effects to each risk gene. In our study, we observed that *Apoe^−/−^* and *Clu^−/−^* mice at the relatively early age of six months show transcriptomic patterns similar to human AD cases. Pathway analysis suggested that *Apoe^−/−^* driven mouse modules specifically affect inflammation/microglia related pathways, while *Clu^−/−^* driven mouse modules have affected neurosignaling, lipid transport, and endocytosis related pathways. These findings suggest that *APOE* and *CLU* risk genes are associated with distinct AD-related pathways. We have also identified that 22 genes were co-expressed in the *Apoe^−/−^*-driven ivory mouse module and in AMP-AD modules from all human brain regions in Consensus Cluster B that were enriched in inflammation and microglia associated pathways. Further, some of these genes *(Tyrobp, Trem2,* and *Csf1r)* were differentially expressed in *Apoe^−/−^* mice. Previous studies have already implicated the role of *TREM2* in AD susceptibility due to association of heterozygous rare variants in *TREM2* with elevated risk of AD [51] and higher cortical *TREM2* RNA expression with increased amyloid pathology [52]. *TYROBP* has been also previously reported as key regulator of immune/microglia associated pathways, which is strongly associated with LOAD pathology [14]. These genes have been also proposed as potential drug targets (https://agora.ampadportal.org/) and our findings supports the role of these genes with pathophysiology of LOAD.

Correlation analysis also identified that mice carrying different mutations capture distinct transcriptional signatures of human LOAD. Moreover, we have observed contrasting correlations of *APOEε4, Apoe^−/−^,* and *Clu^−/−^* mice with AMP-AD modules, implicating that these genetic perturbations might affect LOAD risk through different physiological pathways. It has been speculated that absence of both *Apoe* and *Clu* resulted in accelerated disease onset, and more extensive amyloid deposition in the *PDAPP* transgenic mice brain [53]. Furthermore, APOE and CLU proteins interact with amyloidbeta (Aβ) and regulates its clearance from brain. In particular, the presence of CLU and the *APOEε2* allele promotes Aβ clearance from brain, whereas *APOEε4* reduces the clearance process [43]. These observations also suggest a protective role of *CLU* [43, 54, 55], consistent with our transcriptome-based anti-correlation of *Clu’^−/−^* mice LOAD modules (Figure 6). Understanding of the complex interaction between these genes is essential to interpret molecular mechanisms underlying AD. Hence, it would be interesting to analyze mice models carrying different combinations of genetic variants.

We did not observe any striking responses in brain gene expression patterns in *APOEε4, Bin1^+/−^,* and *Cd2ap^+/−^* mice based on the small subset of differentially expressed genes, as opposed to effects observed in the *Clu^−/−^* and *Apoe^−/−^* models (Table 2). Nor did we observe any mouse modules significantly driven by these perturbations alone. We note that these models were limited to heterozygous mutations in *Bin1* and *Cd2ap* and astrocyte-specific expression of *APOEε4.* The latter limitation may be insufficient to capture the role of *APOE* variants in microglia and disease risk [56]. However, our human-mouse comparison revealed significant correlation of these mouse models with multiple human-derived AMP-AD co-expression modules. We interpret this as these models expression global changes relevant to human cases, while few individual gene expression changes are large enough to be captured by differential expression analysis. This may suggest region-specific and/or cell-specific signals that are diluted by our bulk whole-brain analysis. We have observed that *Bin1^+/−^* models were significantly associated with multiple AMP-AD co-expression modules, which in turn were enriched in immune response, inflammation, and synaptic functioning pathways, which is in concordance with other studies [57, 58]. Furthermore, *Cd2ap^+/−^* mice captured similar human AD signatures as *Clu^−/−^* mice, it may be due to their involvement in similar pathways like blood-brain carrier, and loss of function in *Cd2ap* may contribute to genetic risk of AD by facilitating age related blood-brain barrier breakdown [59]. In-depth investigation of the functional variants of these high-risk AD genes will be essential to evaluate their role in LOAD onset and progression.

The molecular mechanisms of AD driven by rare mutations in *APP*, *PSEN1,* and *PSEN2* are relatively well understood, but the functional impact of LOAD associated risk factors still remain unclear. Although early-onset models have provided critical insights into amyloid accumulation, pathology, and clearance, they do not reflect the full transcriptomic signatures and complete neuropathology of LOAD. Indeed, the primary transcriptomic signatures from mice carrying major early-onset and late-onset genetic factors are distinct (Figure 1B), although our functional analysis in the context of human disease modules also detected some common neuroimmune effects (Figure 6). It remains unclear whether the relatively uncommon EOAD cases and the more common late-onset AD cases proceed through similar disease mechanisms. Understanding these distinctions motivates the development and characterization of new models for the late onset of AD. In this study, we have analyzed mice carrying alterations in LOAD candidate genes and found that different AD risk genes are associated with different AD-related pathways. Our approach provides a platform for further exploration into the causes and progression of LOAD by assessing animal models at different ages and/or with different combinations of LOAD risk variants. This study highlighted that implementing state-of-the-art approaches to generate and characterize LOAD-associated mouse models might be helpful to identify variants and pathways to understand complete AD mechanisms and ultimately develop effective therapies for AD.

## LIST OF ABBREVIATIONS

AD: Alzheimer’s disease
LOAD: Late-onset Alzheimer’s disease
B6: C57BL/6J
RNA-Seq: RNA sequencing
AMP-AD: Accelerating Medicines Partnership for Alzheimer’s Disease
ROSMAP: Religious Orders Study/Memory and Aging Project

## DECLARATIONS

## Acknowledgements

We thank the many institutions and their staff that provided support for this study and who were involved in this collaboration. We would like to acknowledge Ben logsdon for curating human brain data.

## Availability of data and materials

The results published here are in whole or in part based on data obtained from the AMP-AD Knowledge Portal (doi:10.7303/syn2580853). ROSMAP Study data were provided by the Rush Alzheimer’s Disease Center, Rush University Medical Center, Chicago. Data collection was supported through funding by NIA grants P30AG10161, R01AG15819, R01AG17917, R01AG30146, R01AG36836, U01AG32984, U01AG46152, the Illinois Department of Public Health, and the Translational Genomics Research Institute. Mayo RNA-Seq Study data were provided by the following sources: The Mayo Clinic Alzheimer’s Disease Genetic Studies, led by Dr. Nilufer Ertekin-Taner and Dr. Steven G. Younkin, Mayo Clinic, Jacksonville, FL using samples from the Mayo Clinic Study of Aging, the Mayo Clinic Alzheimer’s Disease Research Center, and the Mayo Clinic Brain Bank. Data collection was supported through funding by NIA grants P50 AG016574, R01 AG032990, U01 AG046139, R01 AG018023, U01 AG006576, U01 AG006786, R01 AG025711, R01 AG017216, R01 AG003949, NINDS grant R01 NS080820, CurePSP Foundation, and support from Mayo Foundation. Study data includes samples collected through the Sun Health Research Institute Brain and Body Donation Program of Sun City, Arizona. The Brain and Body Donation Program is supported by the National Institute of Neurological Disorders and Stroke (U24 NS072026 National Brain and Tissue Resource for Parkinson’s Disease and Related Disorders), the National Institute on Aging (P30 AG19610 Arizona Alzheimer’s Disease CoreCenter), the Arizona Department of Health Services (contract 211002, Arizona Alzheimer’s Research Center), the Arizona Biomedical Research Commission (contracts 4001, 0011, 05-901 and 1001 to the Arizona Parkinson’s Disease Consortium) and the Michael J. Fox Foundation for Parkinson’s Research. MSBB data were generated from postmortem brain tissue collected through the Mount Sinai VA Medical Center Brain Bank and were provided by Dr. Eric Schadt from Mount Sinai School of Medicine Mouse RNA-Seq data from the MODEL-AD consortium is available through Synapse via the AMP-AD knowledge portal (www. synapse. org/#! Synapse: syn 15811463)

## Authors’ contributions

RSP performed the gene expression analyses in human and mouse brain tissue. AU and CP supervised the analyses. LG, and GRH performed mouse experiments. GWC and GRH supervised and designed the project. RSP, GWC and CP wrote the manuscript. All authors read and approved the final manuscript.

## Ethics approval

All experiments involving mice were conducted in accordance with policies and procedures described in the Guide for the Care and Use of Laboratory Animals of the National Institutes of Health and were approved by the Institutional Animal Care and Use Committee at The Jackson Laboratory.

## Consent for publication

All authors have approved of the manuscript and agree with its submission.

## SUPPLEMENTAL MATERIAL

**SUPPLEMENTAL TABLE 1: Differentially expressed genes in LOAD mouse strains**

The attached table depicts the differentially expressed genes in LOAD mouse strains compared to C7BL/6J mice.

**SUPPLEMENTAL TABLE 2: KEGG pathway annotation in LOAD mouse strains**

The attached table depicts the KEGG pathway annotations for the up and downregulated genes in LOAD mouse strains compared to C7BL/6J mice.

**SUPPLEMENTAL TABLE 3: Mouse modules of co-expressed genes**

Summaries of the 26 mouse modules of co-expressed genes. In sheet 1, genes in each mouse modules were listed. Sheet 2 depicts mouse modules were observed to be significantly (p < 0.05) driven by at-least one of the mouse strains. Sheets 3 and 4 illustrate enriched KEGG pathways and enriched GO terms in each mouse modules.

**SUPPLEMENTAL TABLE 4: Jaccard indices between Mouse and AMP-AD modules**

The attached table contains Jaccard indices and its significance (p-value) for each mouse-human module pair.

**SUPPLEMENTAL TABLE 5: Transcriptional factor annotations in LOAD mouse modules**

The attached table illustrate transcriptional factor enriched in each 26 mouse modules of co-expressed genes.

